# First-sight recognition of touched objects shows that chicks can solve the Molyneux’s problem

**DOI:** 10.1101/2022.08.18.504388

**Authors:** Elisabetta Versace, Laura Freeland, Shuge Wang, Michael G. Emmerson

## Abstract

If a congenitally blind person learns to distinguish between a cube and a sphere by touch, would they immediately recognise these objects by sight if their vision is restored? This question, posed by Molyneux in 1688, has puzzled philosophers and scientists. We overcame ethical and practical difficulties in the study of cross-modal recognition by using inexperienced chicks. We hatched chicks in darkness, exposed them to tactile smooth vs. bumpy stimuli in darkness and then tested them in a visual recognition task. At first sight, chicks previously exposed to smooth stimuli approached the smooth stimulus significantly more than those exposed to the bumpy stimuli. This shows that cross-modal recognition does not require multimodal experience. At least in precocial species, supra-modal brain areas already function at birth.

---

On 7^th^ July 1688, the philosopher William Molyneux posed a question to John Locke that remains to be addressed: If a person born blind is taught to distinguish between a cube and a sphere by touch, would they then be able to immediately distinguish these objects by mere sight, if their vision is restored (*1*, *2*)? In other words, is learning required to associate tactile (proprioceptive) and visual (extrapersonal) sensory information that are mediated by different sensory modalities (*3*), or do we automatically build internal representations of objects that share supra-modal or multimodal features (*4*)?

To solve the Molyneux’s question and understand whether cross-modal recognition exists in the absence of cross-modal experience, we need to exclude a role of previous experience. To this end, it is necessary to overcome ethical problems regarding limitation of experience and practical issues on the functionality of sensory systems. When sight is restored after long-term blindness or in congenitally blind patients, for instance via surgical removal of the cataract, sight can be impaired. For this reason, the lack of transfer from tactile discrimination to vision observed immediately after the onset of sight (from Cheseldon 1798, to the more recent ones (*5*)) can be due to misperception (*2*) or degradation of the visual system (*6*). On the contrary, successful transfers can be due to learning from verbal descriptions or associations from other sensory modalities. Another approach to the puzzle has focused on early life responses. Tactual-visual recognition has been found in human newborns exposed to pacifiers with different shapes (*7*). Yet, this early performance could not rule out the effect of one month of previous experience. In humans, it is not ethical to conduct studies where subjects are sensorily deprived and cannot experience any association between stimuli from specific sensory modalities before the test.

These problems can be addressed studying precocial species such as domestic chicks (*Gallus gallus*), that are an ideal model to understand cognition at the beginning of life (*8*–*10*). First, chicks hatch with already mature visual, proprioceptive and motor systems, so that their perceptive and motor responses can be evaluated in the first hours of life (*8*, *10*, *11*). Second, chicks’ vision is not impaired by temporary deprivation of visual experience (*10*, *11*). Furthermore, investigating cross-modal recognition in non-human animals rules out any influence of verbal reports.

Our experiment comprised three phases: hatching, exposure and test. We hatched chicks in individual compartments in darkness. Hatching boxes contained either smooth cubes or bumpy cubes (Fig. 1A-B, Fig. S1). In the smooth condition, bumps were located in a separate subcompartment. Chicks remained for 24 hours in their hatching box, where they could explore the smooth or bumpy stimuli through their tactile sensory modality. Infrared camera recordings showed that chicks hatched in darkness explore their surroundings (10% time moving, Fig. 1C) and spend a large part of their time in contact with the experimental objects (60% of time in contact with stimuli, Fig. 1D). This is not surprising because, in the wild, chicks spend the first three days of life mainly under the mother hen (*12*). Hence, movement and tactile exploration in darkness is a common experience for newly-hatched chicks. After tactile exposure, chicks were individually moved to the testing room in an opaque box and presented with a visual recognition task of the smooth vs. bumpy stimuli. Test stimuli (that were identical to stimuli used during dark-reared stimulus exposure, but not those actually used during exposure) were located on slowly rotating platforms, to enhance approach responses in chicks. Chicks could freely move in the arena, while a wire mesh prevented any tactile interactions with the stimuli (Fig. 1B), see Materials and Methods for details.

**Fig. 1.**
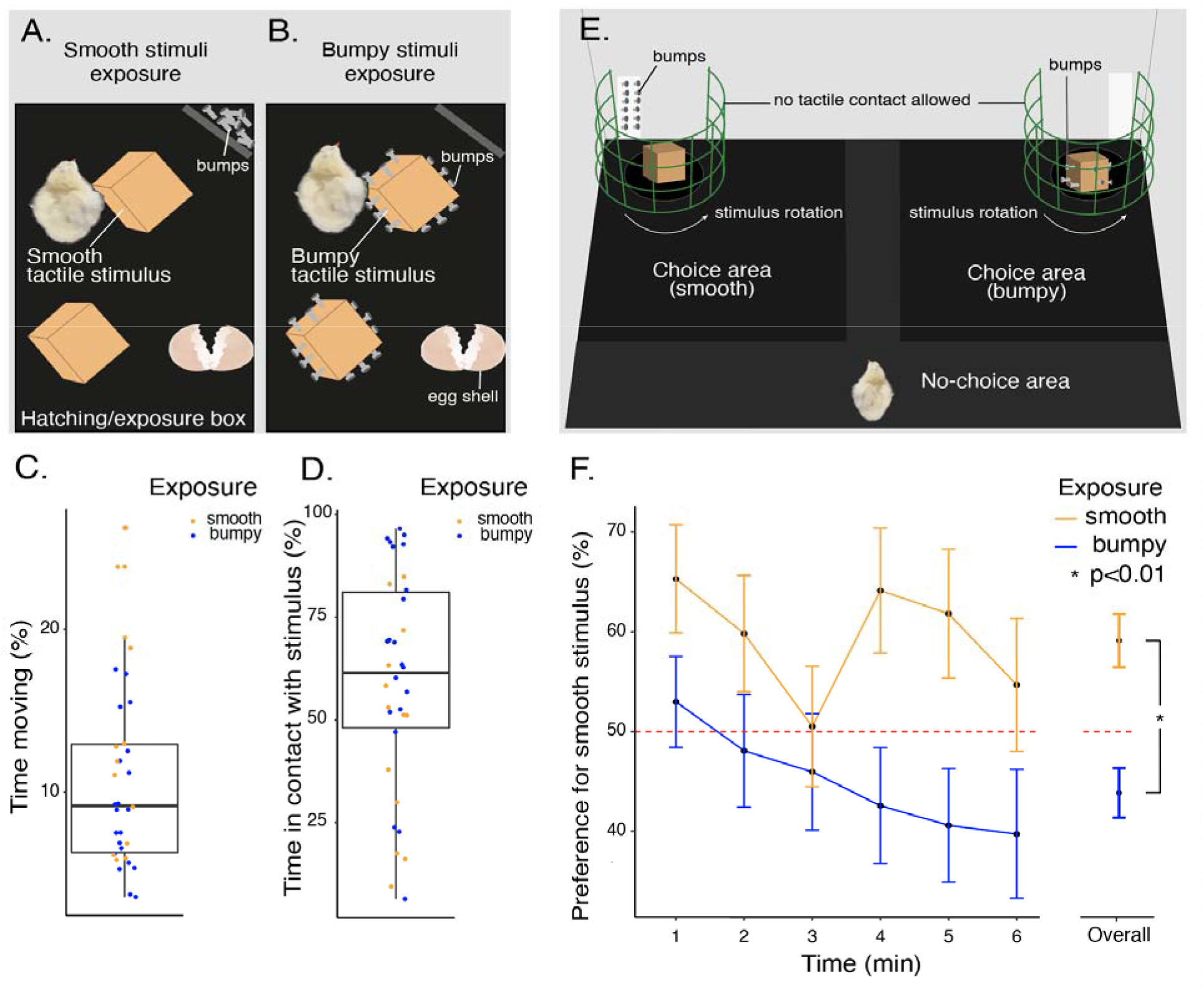
Experimental setting and results. Hatching/exposure compartments contained the smooth (**A**) and bumpy (**B**) stimuli that chicks experienced in darkness for 24 hours. In darkness, newly hatched chicks spend about 10% of the time moving (**C**) and about 50% of their time in touch with the tactile stimuli we provided (**D**). After exposure, chicks were moved to the testing arena (**E**), where they could visually explore and approach the smooth and bumpy stimulus. The test stimuli were located behind a mesh and could not be touched. The right-left position of the stimuli was counterbalanced between chicks. For visual and olfactory matching, bumps (metal bolts) were present on both sides of the arena. The (**F**) panel shows the preference for the smooth stimulus (Mean +/− SEM) in chicks previously exposed to the smooth (orange) and bumpy (blue) stimuli. The overall preference across the six minutes of observation is shown on the right side.

We hypothesised that, during the exposure phase, chicks use tactile experience to learn the smooth vs. bumpy tactile quality of the stimuli, similarly to what happens in visual filial imprinting. Filial imprinting is a learning mechanism based on exposure, where young animals become attached to the objects they are exposed to, without any reinforcement. As a result, after imprinting exposure, chicks tend to approach familiar objects (*13*–*16*). This phenomenon has been widely documented in visual and acoustic modality (*17*–*19*). Although chicks spend most of their first days in contact with the hen (*12*), it is not known whether imprinting works in tactile modality. However, tactile sensory experience is very important for young birds: in several precocial species, visual preferences for objects are enhanced when birds can also touch the objects (*20*, *21*). Our experiment can clarify whether tactile experience can affect chicks’ visual preferences.

In our setting, after tactile exposure in darkness, chicks immediately experienced the visual recognition test. In their first six minutes of visual experience, chicks exposed to the smooth stimuli spent more time close to the smooth stimuli than chicks exposed to the bumpy stimuli (F_1,69_=7.178, p=0.009, Fig. 1E, Table S1). No significant differences between sexes and time points or interactions have been detected. These results show that visually-naïve chicks learn about objects presented in the solely tactile modality, and can use representations based on tactile experience to solve a visual recognition task. This evidence suggests that supra-modal brain areas (*4*, *22*) might be already functional at birth, at least in precocial species, without the need of experience and multi-modal associations. Cross-modal visuo-tactile recognition is likely mediated by tactile imprinting.

The Empiricist John Locke and his followers thought that the Molyneux’s question cannot be solved without previous experience (*23*). Newly-hatched chicks contradict this claim, showing that the brain is equipped to spontaneously match visual and tactile information at birth.

## Materials and Methods

### 1) Cross modal recognition experiment

#### Subjects, incubation, hatching

Overall, we tested 103 chicks (55 females, 48 males) (*Gallus gallus*) of the Ross 308 strain, and 73 chicks (40 females, 33 males) made a choice during the test and were analysed. Fresh eggs were collected from Avara Foods (Rayne, UK), kept at 11-12° C before incubation, incubated in darkness at 37.7° C and 40-60% humidity in a Fiem incubator (model MG 140) for 18 days. Incubation and hatching happened in complete darkness. The incubator and hatchery were located in a room with no windows and with a double blackout curtain system at the door, to prevent any light filtering. The ventilation openings of incubators and hatcheries were covered with blackout fabric. At day 18 of incubation, eggs were moved to individual hatching compartments (28×18×11 cm) in a Fiem hatchery (MG316). Each hatching compartment contained either two smooth or two bumpy stimuli (Fig. 1A-B). Each hatchery contained up to 12 hatching compartments, stacked in three layers.

**Fig. S1.**
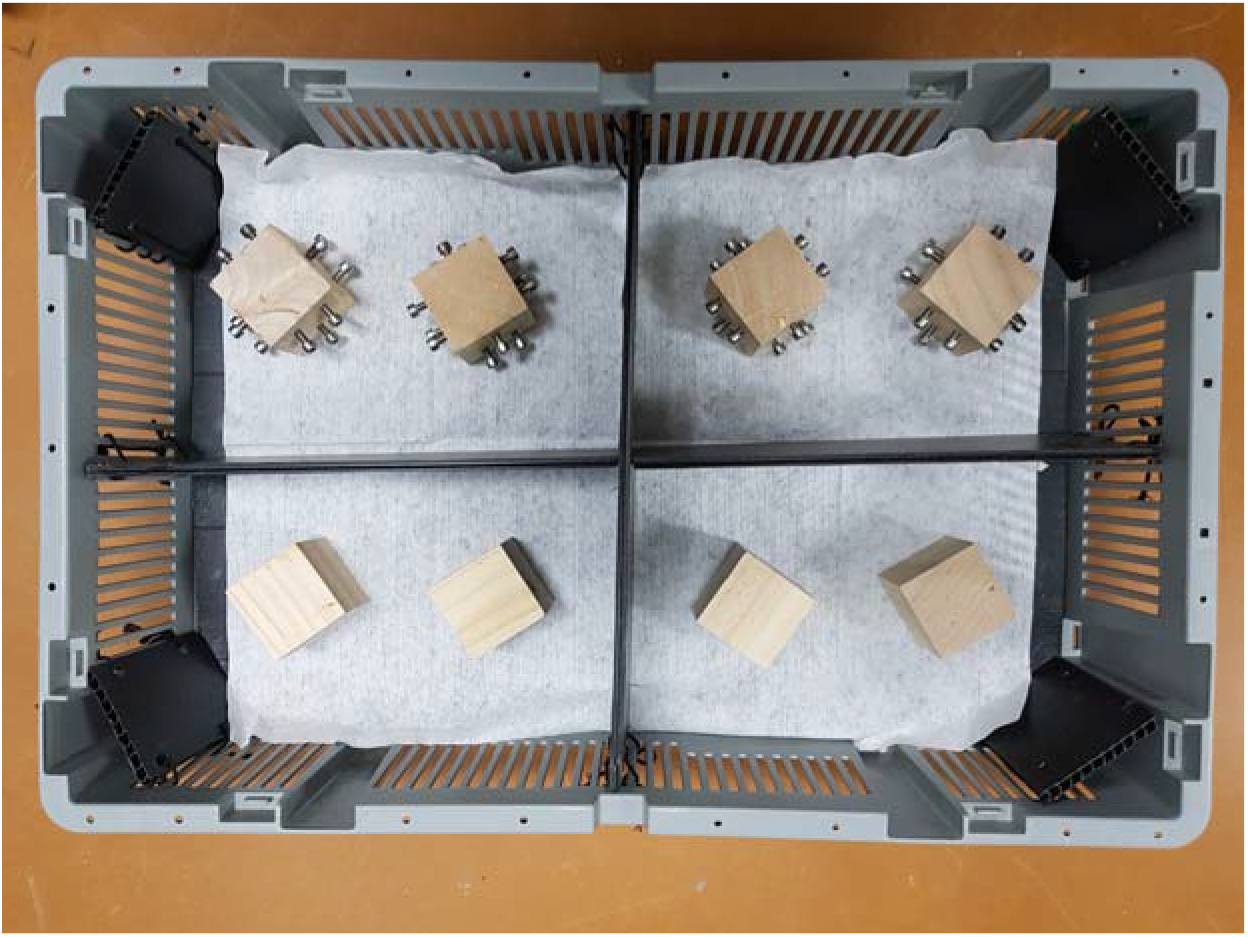
Picture of a hatching tray containing four hatching compartments. Hatching trays were gray plastic trays with vertical gaps for ventilation and split into four equally sized hatching compartments using black plastic secured to the exterior walls with strings. Each hatching compartment base was lined with a paper towel and contained two objects for tactile exposure. Two bumpy objects and two smooth objects are shown in the top and bottom part of the figure respectively. Each hatching compartment contained a sub-compartment with either 12 bolts (smooth condition) or nothing (bumpy condition) to minimise differences between conditions.

#### Stimuli

Tactile stimuli were wooden cubes (5 cm side), with or without metal bolts (16 mm stainless steel) drilled on the sides as bumps. Bumpy stimuli had three bolts embedded into their four side walls (12 bolts total), protruding 10 mm (see Figure 1B). Smooth stimuli had smooth sides, and 12 bolts were located in a separate subsection of the compartment. Previous research showed that metal doesn’t smell (*24*), but to equalize the quantity of material located throughout the compartment, we created a separate section for the bolts in the smooth condition. This subsection was left empty in the bumpy condition.

#### Apparatus

The test apparatus is displayed in Fig. 1E, Fig. S2. The apparatus was a 90 (wide) x 60 (long) x 52 (tall) cm arena with white walls and a black non-slip mat floor. Two PrimeMatik electric black rotary platforms (15.1 cm diameter), moving at 2.4 rotations/minute were located at the top right and left regions of the apparatus. The stimulus was located approximately 0.5 cm above the floor of the arena. The tactile stimuli were placed on the rotators (Fig. 1C). The rotary platforms rotated in the same direction (i.e., clockwise or anticlockwise). Green plastic mesh surrounded each rotator, preventing the chick from interacting with the stimuli. On the corner behind the rotators, a plastic box was left empty (behind the smooth stimulus) or filled with 12 bolts drilled (smooth stimulus). This controlled potential differences cues between conditions. An LED stripe above the apparatus illuminated the arena. Experimental sessions were recorded with a Microsoft LifeCam Studio webcam located in the centre of the arena. For analysing chicks’ preferences, the arena was virtually divided in three areas: a left area (a 40×40 cm square close to one stimulus), a right area (a 40×40 cm rectangle close to the other stimulus) and a no-choice area (see Fig. S2).

**Fig. S2.**
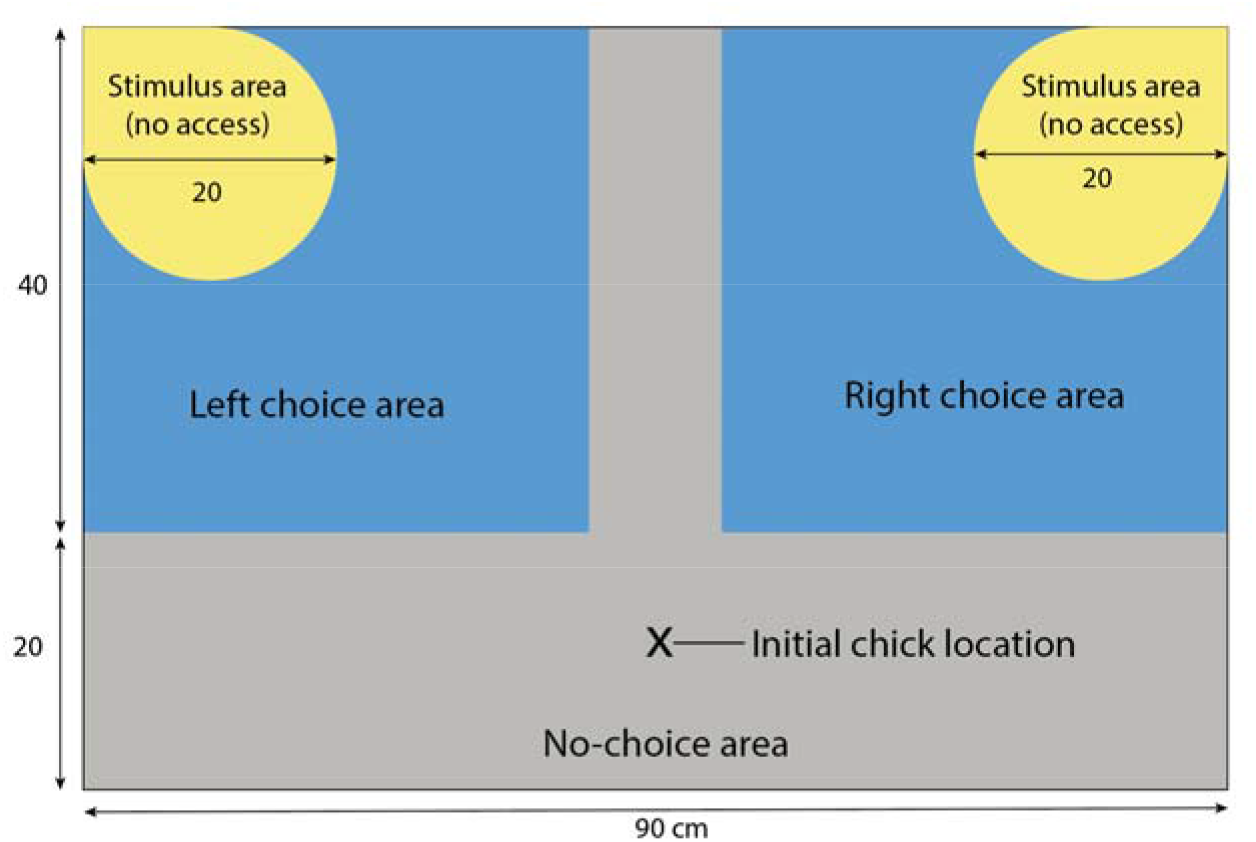
Schematic diagram of the testing arena used for the visual-recognition test. The arena was composed of two stimulus areas not accessible to chicks (illustrated in yellow), a left and right choice area (illustrated blue), and a no-choice area (illustrated in grey). Stimulus areas contained the rotators and stimuli, enclosed behind a mesh barrier.

#### Procedure

Upon hatching, each chick could move around within its compartment and touch the two identical tactile stimuli (Fig. 1A-B). In darkness, experimenters manually checked that chicks hatched by 10 AM and were healthy. The exposure phase lasted 24-30 hours, after which chicks were individually tested. Each chick was moved into an individual opaque box, covered with a lid, and gently transported to the testing room. At test, each chick was sexed and gently located in the testing arena, in the no-choice area opposite to the stimuli (Fig. 1E and S2). Chicks were observed and their performance analysed for 6 minutes. Chicks that didn’t make a choice within 6 minutes were not included in statistical analyses, in line with previous literature (*17*, *25*–*27*).

#### Video and data analysis

The position of the centroid of the chick in the arena has been analysed with the open source automated behavioural tracking system DeepLabCut (*28*). For the centroid of each chick, we ensured that the quality of tracking reached an accuracy of 0.9 likelihood for at least 90% of frames, since this threshold ensured high accuracy.

For each of the six 1-minute time bins, the Preference for the smooth stimulus has been calculated as: (Time spent in the smooth stimulus area)/(Time spent in the smooth stimulus area + Time spent in the bumpy stimulus area)*100

We used R (*29*) (packages: ggplot2, rstatix, plyr) for statistical analysis and plots.

We ran a mixed design ANOVA with Preference for smooth stimulus as dependent variable, Exposure condition (smooth or bumpy) and Sex (female or male) as between subjects variables and Time bin (1-6) as within-subjects variable. Alpha was set to p<0.05.

#### Results

The full ANOVA results are presented in Table S1.

**Table S1.**
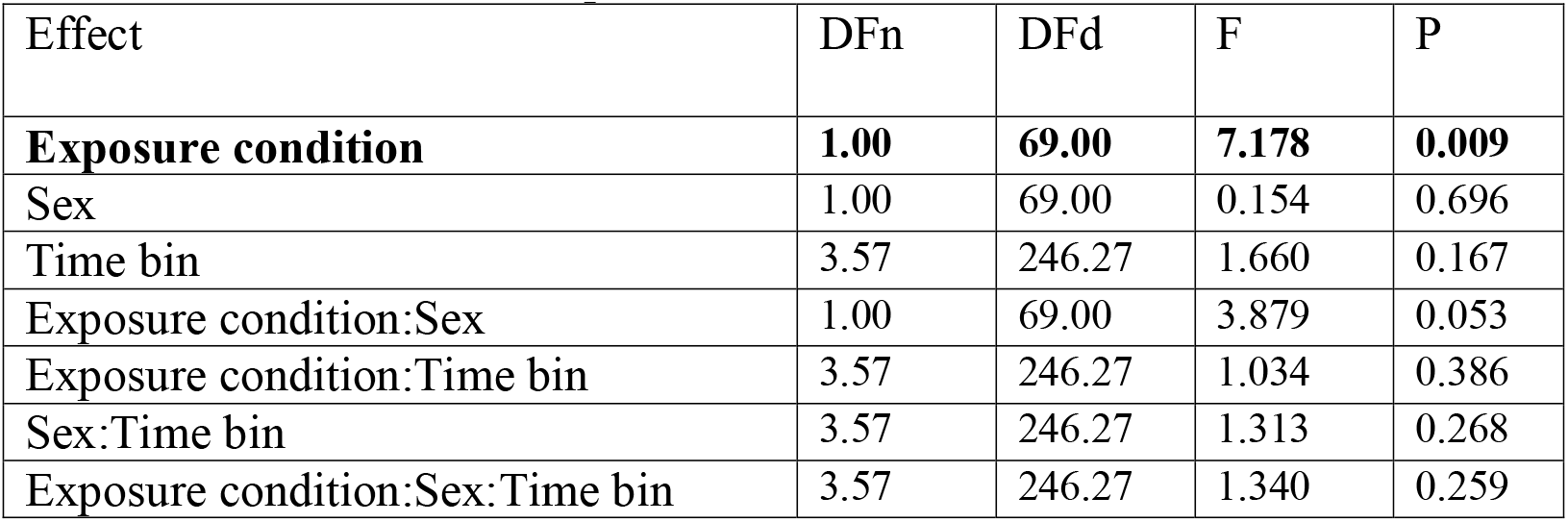
Output table from an ANOVA, displaying all main effects and interactions. Bold indicates significant effects.

### 2. Monitoring tactile stimulation in the hatching compartments

#### Subjects, incubation, hatching, stimuli

We tested 34 chicks (15 females, 19 males) (*Gallus gallus*) of the Ross 308 strain. Fresh eggs were collected from the Avara Food hatchery (Rayne, UK), maintained, incubated and hatched in the same conditions and with the same stimuli described in the previous experiment.

#### Apparatus

Since the hatcheries where too small to host infrared cameras, recordings of the activity of chicks during exposure to the tactile stimuli were conducted on different chicks from those tested in the previous experiment, in a pre-warmed (30-32° C) room, in full darkness. A double door system prevented any light filtering in the room. The same hatching/exposure compartments described in the previous experiment were used. We recorded chicks’ movement using a Foscam C2M infrared camera.

#### Procedure

After hatching in the tray with tactile stimuli, chicks were individually moved from the hatchery to an identical compartment located in a pre-warmed room. For the transport, chicks were located in individual opaque boxes in darkness, and boxes were located in an opaque bag.

Chick were then gently located in a hatching/exposure tray and recorded for 24 hours.

#### Video analysis

For the analysis of chicks’ position, we virtually divided the rectangular area of the exposure comportment in 6 squares (a-f, see Fig. S3). We assessed chicks’ behaviour for 5 minutes every hour (2 out 24 hours of recording), sampling at 20 second intervals (15 sampling times) in each hour. We recorded the position (a-f) of the centroid of the chick and whether the chicks were touching (or not) one tactile stimulus.

**Fig. S3.**
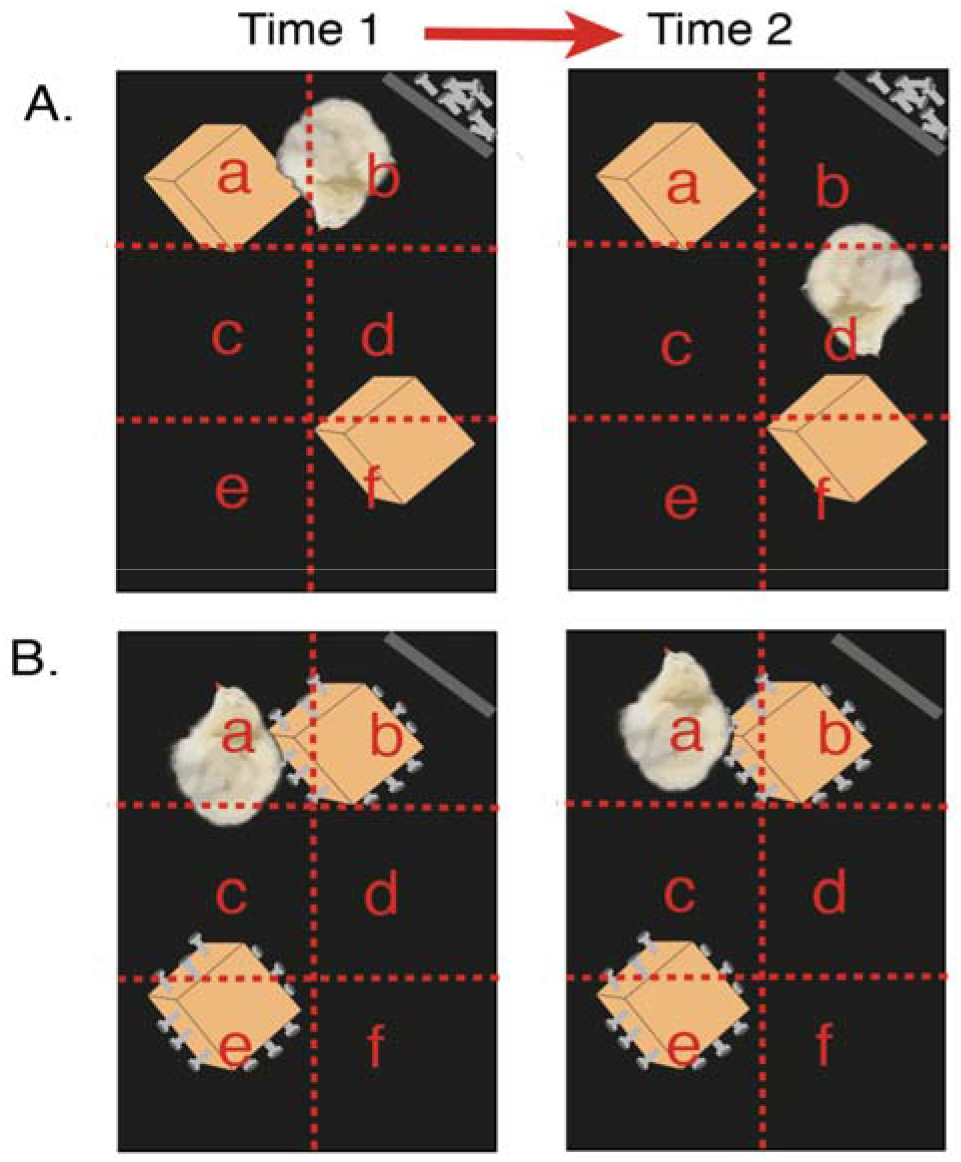
Diagram showing how a chick’s behaviour was quantified during tactile exposure monitoring in the hatching compartment. Each compartment was divided into six equally sized areas (labelled a-f). Scan sampling was used to measure the percentage of time in which a chick was moving (centroid located in two different areas between two subsequent time points) and the percentage of time in which chicks were in touch with the tactile stimulus. In panel (**A**), the chick is in contact with the stimulus at Time 1 but not at Time 2, and is moving between Time 1 and Time 2. In panel (**B**), a chick is contact with the stimulus in Time 1 and Time 2 and is not moving between Time 1 and Time 2.

#### Data analysis and results

We estimated the average percentage of time that each chick spent moving as: (Number of observations in which the chick’s centroid changed position)/(Total number of observations)*100.

Overall (Fig. 1C), chicks spent on average 10.78% (SD=5.65) of their time moving (median 9.17%); Smooth object condition: average=13.15%, SD=6.91, median=11.90%; Bumpy object condition: average=9.33%, SD=4.26, median=11.90. This shows that chicks did move in darkness.

We estimated the average percentage of time that each chick spent in contact with the tactile stimulus as: (number of observations in which the chick touched the stimulus)/(total number of observations)*100.

Overall (Fig. 1D), chicks spent on average 59.07% of their time (median 61.51%, SD=26.45), namely >14 hours, in touch with the tactile stimulus; Smooth object condition: mean=48.28% (about 12 hours), SD=24.84, median=51.21%; Bumpy object condition: mean=65.75%, (>15 hours), SD=25.73, median=68.89. These results show that in all conditions chicks had plenty of direct exposure time to imprint in tactile modality.

## Ethics Statement

The experiments complied with national laws on the use of animals in research and were approved by the Home Office (PPL number: PP5180959) and by the Queen Mary University of London Animal Welfare and Ethical Review Body.

## Acknowledgments

We thank the Prepared Minds (Versace) lab and Chittka lab members at Queen Mary University of London for fruitful discussions on the project. We thank Kiera Rose for help with a related pilot study. We thank Avara Foods that generously donated the eggs for these experiments.

## Author contributions

Conceptualization: EV; Methodology: EV, LF, ME, SW; Investigation: LF, SW, ME; Visualization: EV; Funding acquisition: EV; Project administration: EV, LF; Supervision: EV; Writing – original draft: EV; Writing – review & editing: EV, LF, SW, ME

## References

1. D. Marjolein, G.-J. Lokhorst, “Molyneux’s Problem.” Stanford Encycl. Philos. (2021), (available at https://plato.stanford.edu/archives/win2021/entries/molyneux-problem/).

2. M. Degenaar, Molyneux’s problem: three centuries of discussion on the perception of forms (Kluwer Academic Publishers, Dordrecht, Netherlands, 1996).

3. B. Glenney, Adam Smith and the Problem of the External World. J. Scottish Philos. 9, 205–223 (2011).

4. X. Wang, W. Men, J. Gao, A. Caramazza, Y. Bi, Two Forms of Knowledge Representations in the Human Brain. Neuron. 107, 383–393.e5 (2020).

5. R. Held, Y. Ostrovsky, B. deGelder, P. Sinha, Revisiting the molyneux question. J. Vis. 8, 523 (2008).

6. I. Fine, A. R. Wade, A. A. Brewer, M. G. May, D. F. Goodman, G. M. Boynton, B. A. Wandell, D. I. A. MacLeod, Long-term deprivation affects visual perception and cortex. Nat. Neurosci. 6, 915–916 (2003).

7. A. N. Meltzoff, R. W. Borton, Intermodal matching by human neonates. Nature. 282, 403–404 (1979).

8. E. Versace, A. Martinho-Truswell, A. Kacelnik, G. Vallortigara, Priors in Animal and Artificial Intelligence: Where Does Learning Begin? Trends Cogn. Sci. 22, 963–925 (2018).

9. E. Versace, G. Vallortigara, Origins of knowledge: Insights from precocial species. Front. Behav. Neurosci. 9 (2015), doi:10.3389/fnbeh.2015.00338.

10. O. Rosa-Salva, U. Mayer, E. Versace, H. Marie, B. S. Lemaire, G. Vallortigara, Sensitive periods for social development: Interactions between predisposed and learned mechanisms. Cognition. 213, 104552 (2021).

11. E. Versace, G. Vallortigara, Origins of knowledge: Insights from precocial species. Front. Behav. Neurosci. 9, 338 (2015).

12. C. J. Nicol, The Behavioural Biology of Chickens (CABI, Oxfordshire OX10 8DE (UK), 2015).

13. B. J. McCabe, Visual Imprinting in Birds: Behavior, Models, and Neural Mechanisms. Front. Physiol. 10 (2019), doi:10.3389/fphys.2019.00658.

14. P. Bateson, The characteristics and context of imprinting. Biol. Rev. 41, 177–217 (1966).

15. K. Lorenz, The Companion in the Bird’s World. Auk. 54, 245–273 (1937).

16. J. J. Bolhuis, Mechanisms of avian imprinting: a review. Biol. Rev. 66, 303–345 (1991).

17. E. Versace, M. J. Spierings, M. Caffini, C. ten Cate, G. Vallortigara, Spontaneous generalization of abstract multimodal patterns in young domestic chicks. Anim. Cogn. 20, 521–529 (2017).

18. G. Batista, J. L. Johnson, E. Dominguez, M. Costa-Mattioli, J. L. Pena, Regulation of filial imprinting and structural plasticity by mTORC1 in newborn chickens, 1–11 (2018).

19. G. Batista, J. L. Johnson, E. Dominguez, M. Costa-Mattioli, J. L. Pena, Translational control of auditory imprinting and structural plasticity by eIF2α. Elife. 5, 1–19 (2016).

20. M. Clements, J. Lien, Effects of tactile stimulation on the initiation and maintenance of the following response in Japanese quail *(Coturnix coturnix japonica)*. Anim. Learn. Behav. 3, 301–304 (1975).

21. L. A. Eiserer, The effects of tactile stimulation on imprinting in ducklings after the sensitive period. Anim. Learn. Behav. 6, 27–29 (1978).

22. E. Ricciardi, D. Bonino, S. Pellegrini, P. Pietrini, Mind the blind brain to understand the sighted one! Is there a supramodal cortical functional architecture? Neurosci. Biobehav. Rev. 41, 64–77 (2014).

23. G. Ferretti, B. Glenney, Molyneux’s queston and the history of philosophy (Routledge, London, 2020).

24. D. Glindemann, A. Dietrich, H. J. Staerk, P. Kuschk, The two odors of iron when touched or pickled: (Skin) carbonyl compounds and organophosphines. Angew. Chemie - Int. Ed. 45, 7006–7009 (2006).

25. L. Regolin, L. Tommasi, G. Vallortigara, Visual perception of biological motion in newly hatched chicks as revealed by an imprinting procedure. Anim. Cogn. 3, 53–60 (2000).

26. M. Hernik, A. Broseghini, G. Vallortigara, Visually-naïve chicks prefer agents that move as if constrained by a bilateral. Cognition. 173, 106–114 (2018).

27. E. Versace, J. Schill, A. M. Nencini, G. Vallortigara, Naïve chicks prefer hollow objects. PLoS One. 11 (2016), doi:10.1371/journal.pone.0166425.

28. A. Mathis, P. Mamidanna, K. M. Cury, T. Abe, V. N. Murthy, M. W. Mathis, M. Bethge, DeepLabCut: markerless pose estimation of user-defined body parts with deep learning. Nat. Neurosci. 21, 1281–1289 (2018).

29. R. C. Team, A language and environment for statistical computing. (2022), (available at https://www.r-project.org/).

